# Inheritance of DNA methylation differences in the mangrove Rhizophora mangle

**DOI:** 10.1101/2020.10.24.353482

**Authors:** Jeannie Mounger, M. Teresa Boquete, Marc W. Schmid, Renan Granado, Marta H. Robertson, Sandy A. Voors, Kristen L. Langanke, Mariano Alvarez, Cornelis A.M. Wagemaker, Aaron W. Schrey, Gordon Fox, David B. Lewis, Catarina Fonseca Lira, Christina L. Richards

**Author notes:** shared first author (Jeannie Mounger, M. Teresa Boquete, Marc W. Schmid).

## Abstract

The capacity to respond to environmental challenges ultimately relies on phenotypic variation which manifests from complex interactions of genetic and non-genetic mechanisms through development. While we know something about genetic variation and structure of many species of conservation importance, we know very little about the non-genetic contributions to variation. *Rhizophora mangle* is a foundation species that occurs in coastal estuarine habitats throughout the neotropics where it provides critical ecosystem functions, and is potentially threatened by climate change. Several studies have documented landscape level patterns of genetic variation in this species, but we know virtually nothing about the inheritance of non-genetic variation. To assess one type of non-genetic variation, we examined the patterns of DNA sequence and DNA methylation in maternal plants and offspring from natural populations of *R. mangle* from the Gulf Coast of Florida. We used a reduced representation bisulfite sequencing approach (epi-genotyping by sequencing or epiGBS) to address the following questions: a) What are the levels of genetic and epigenetic diversity in natural populations of *R. mangle*? b) How are genetic and epigenetic variation structured within and among populations? c) How faithfully is epigenetic variation inherited? We found low genetic diversity but high epigenetic diversity from natural populations of maternal plants in the field and that a large portion (up to ~25%) of epigenetic differences among offspring grown in common garden was explained by maternal family. Therefore, epigenetic variation could be an important source of response to challenging environments in the genetically depauperate populations of this foundation species.

## 1. Introduction

Preserving the ability of populations to respond to environmental challenges is critical to conservation efforts. This ability ultimately depends on phenotypic variation (Norberg et al., 2001; Björklund et al., 2009; Henn et al., 2018), and consequently conserving genetic variation has been championed by numerous researchers studying conservation in recent decades. However, the focus on genetic variation must be interpreted with caution (Hufford and Mazer, 2003) considering the misplaced emphasis on the concept that only variation in DNA sequence matters (Keller, 2002, 2014; Sultan, 2015; Bonduriansky & Day, 2018). In fact, Sultan (2015) argued that as modern biologists our task is to restore the context dependence of gene expression and trait variation which has become particularly relevant in the context of anthropogenic alterations to natural ecosystems. In the framework of re-evaluating the mapping of genotype to phenotype (Pigliucci, 2010; Keller, 2014), we can now use the concepts of Evo-Devo to explore plasticity and structure within populations, as well as examine how these processes are impacted by climate change (Campbell, Adams, Bean, & Parsons, 2017).

Natural epigenetic variation (e.g., alterations to DNA methylation, small RNAs, and chromatin remodeling) has been associated with phenotypic and functional diversity in plants, emerging both as a molecular-level mechanism underlying phenotypic plasticity and as a potentially important non-genetic source of heritable variation (Medrano, Herrera, & Bazaga, 2014; Cortijo et al., 2014; Balao, Paun, & Alonso, 2018; Banta & Richards, 2018; Zhang, Latzel, Fischer, & Bossdorf, 2018). There is increasing evidence that suggests that environmentally-induced epigenetic variation can be heritable, particularly in plants (e.g. Verhoeven, Jansen, Van Dijk, & Biere, 2010; Richards et al., 2012; Herrera et al., 2017) but this contention is not universally supported (reviewed in Richards & Pigliucci, *in press*). This source of variation may be imperative for sessile organisms as they cope with a broad range of environmental conditions without the ability to migrate away from stressors (Balao, Paun, & Alonso, 2018). Further, rapid phenotypic alterations mediated by epigenetic mechanisms may be especially important for the persistence of plant populations in dynamic ecosystems that endure significant natural and anthropogenic environmental variability, such as those in coastal and alpine regions (Nicotra et al., 2015; Burggren, 2016).

Much of what is presently known about the functionality of epigenetic variation predominantly comes from studies of model organisms (Richards et al., 2017; Balao, Paun, & Alonso, 2018). For instance, epigenetic differences in *Arabidopsis thaliana* have been linked to response to temperature (Kawakatsu et al., 2016) and biotic stressors (Dowen et al., 2012; reviewed in Zogli & Libault, 2017). Additionally, inheritance of epigenetic variation has been observed in *A. thaliana* (Lang-Mladek et al., 2010; Blevins et al., 2014) as well as in several crop species (e.g. DNA methylation in maize and *Fragaria vesca*; Li et al., 2014; de Kort et al., 2020; and small RNAs in *Brassica rapa*; Bilichak et al., 2014). Our understanding of how epigenetic variation behaves in ecological contexts is far more limited, however. Common garden studies of non-model plant species have elucidated changes in DNA methylation that are linked to community composition (van Moorsel et al., 2019) and responses to temperature and nutrient stress (Verhoeven, Jansen, Van Dijk, & Biere, 2010; Nicotra et al., 2015).

Moreover, methylation modifications in natural plant populations are known to be associated with response to habitat and environmental variation (Foust et al., 2016; Gáspár, Bossdorf, & Durka, 2019), hybridization and allopolyploidization (Salmon, Ainouche, & Wendel, 2005; Sehrish et al., 2014; reviewed in Mounger, et al., 2020), fluctuations in salinity and nutrient levels (Lira-Medeiros et al., 2010), light availability (Schulz et al., 2014), and biotic interactions (e.g. herbivory; Herrera & Bazaga, 2011; reviewed in Alonso, Ramos-Cruz, & Becker, 2018). However, other studies have shown that epigenetic variation accumulated following single genetic mutations (Becker et al., 2011; Dubin et al., 2015; Sasaki et al., 2019) and many authors have argued that epigenetic variation is ultimately explained by genetic variation (Alvarez et al., 2020; Robertson et al., 2020).

Understanding the mechanisms of response in coastal foundation species has become increasingly important for conservation and management strategies as these species must cope with rising sea levels and increased warming due to climate change (Osland, Enwright, Day, & Doyle, 2013; Osland et al., 2017b). Worldwide, mangrove forests perform significant ecosystem services including buffering storm surges and tidal wave action, reducing erosion, sequestering an estimated 34.4 Tg of carbon per year (Mcleod et al., 2011), and providing habitat for economically important marine fauna (Alongi, 2008). These forests also play important roles in nutrient and sediment dynamics that are integral to the ecosystem processes of several marine systems, notably coral reefs and seagrass flats (Alongi, 2008; Polidoro et al., 2010). Despite their importance, the distribution and persistence of mangrove tree species are threatened by historic and current land-use change as well as by pollution from agriculture and urban runoff, sewage effluents, hazardous materials spills, and other contaminants from human activities (Ellison, Farnsworth, & Moore, 2015). The Food and Agriculture Organization of the United Nations (FAO) estimates that approximately 35% of global mangrove forest habitat has been destroyed since roughly 1980 for the development of human settlements, agriculture and aquaculture, and industrial shipping harbors (FAO 2007; Polidoro et al., 2010; Ellison et al., 2015). In some regions, mangrove trees are also harvested for wood and charcoal (Ellison et al., 2015), resulting in habitat fragmentation and isolation of existing remnant fragments (Haddad et al., 2015; Friess et al., 2012).

While most mangrove species are not considered to be threatened, 16% of true mangrove species (as defined by Tomlinson, 2016) have qualified for listing on the Red List of Threatened Species of the International Union for Conservation of Nature (IUCN) (Alongi, 2008; Polidoro et al., 2010; Tomlinson, 2016), despite restoration efforts and governmental protections (Lai et al., 2015; Ferreira & Lacerda, 2016). Besides, habitat loss continues to be a serious threat, with current average annual rates of loss of 1-2% (Alongi, 2008; Polidoro et al., 2010). The resultant loss of diversity could pose risks for these coastal foundation species in the future, particularly as sea levels are projected to rise between 0.2 and 2 m over the next century due to anthropogenic climate change (Melillo, Richmond, & Yohe, 2014). Although mangrove tree species may not be at immediate risk of extinction, the degradation and loss of high-quality mangrove forests negatively impacts ecosystem processes, trophic states, and food availability, which in turn threatens biodiversity within these systems (Carugati et al., 2018). On the other hand, evidence has suggested that populations of many mangrove species have historically moved along the intertidal zone and poleward at pace with changes in sea level, reduced incidence of winter frost, and a variety of other abiotic conditions (Alongi, 2008; Osland et al., 2017a). The mechanisms that allow for this migration are not well understood (Osland, Enwright, Day, & Doyle, 2013; Osland et al., 2017b) and coastal development poses a significant barrier to these species’ abilities to colonize landward (Polidoro et al., 2010; Schuerch et al., 2018; reviewed in Godoy & Lacerda, 2015).

To date, broad surveys of genetic diversity across the expansive ranges of mangrove species are lacking, and virtually no studies have directly addressed the importance of non-genetic variation for the persistence of coastal plant species. However, genetic variation has been investigated in limited geographic regions in order to assess patterns of evolution (Duke, Lo, & Sun, 2002), hybridization and introgression (Cerón-Souza et al., 2010), genetic population and subpopulation structure (Arbelaez-Cortes Castillo-Cardenas, Toro-Perea, & Cardenas-Henao, 2007; Ceron-Souza et al., 2010; Albrect Kneeland, Lindroth, & Foster 2013; Bruschi et al., 2014; Chablé Iuit et al., 2020), and to evaluate range expansion as a result of climate change (Sandoval-Castro et al., 2012; Kennedy et al., 2016) in *Rhizophora mangle*.

*Rhizophora mangle* populations appear to vary tremendously in genetic variation across their range. For example, populations along the Pacific coast have greater genetic diversity than those sampled elsewhere within their range across the Western Hemisphere (Arbelaez-Cortes et al., 2007; Cerón-Souza, Bermingham, McMillan, & Jones, 2012; Bruschi et al., 2014). Other studies also suggest that *R. mangle* populations are not panmictic, and instead tend to form somewhat isolated groups (Pil et al., 2011). Populations of *R. mangle* can become genetically isolated both at range ends and in areas of limited tidal flow (Sandoval-Castro et al., 2012; Kennedy et al., 2016), and its poleward expansion is limited by freezing events (its current northern range limit is roughly 29° N latitude; Kennedy et al., 2016). Populations at these peripheries could require particular conservation attention since they have been shown to have greater genetic differences among populations and reduced genetic diversity (Polidoro et al., 2010; Sandoval-Castro et al., 2012; Kennedy et al., 2016), which has been attributed to limits in dispersal ability, low effective population size, a reduction in pollinators, and increased environmental pressures (Sandoval-Castro et al., 2012). In addition, increased warming as a consequence of climate change could result in either the relaxation or amplification of some of these biotic and abiotic limitations at range ends (Devaney, Lehmann, Feller, & Parker, 2017).

In this study, we used the reduced representation bisulfite sequencing approach epigenotyping by sequencing (epiGBS; van Gurp et al., 2016) to measure genetic and DNA methylation differentiation among red mangrove populations near the northern limit of this species in the Tampa Bay region. We took advantage of the unusual biology of *R. mangle* that allows for collecting viviparous propagules that are still attached to the maternal plant. From six populations we collected leaves from maternal trees and their offspring propagules to answer the following questions: a) What are the levels of genetic and epigenetic diversity in natural populations of *R. mangle*? b) Are genetic and epigenetic variation structured among populations of this species in the wild? c) To what extent does epigenetic variation in the offspring correlate with the maternal plants?

## 2. Materials and Methods

### 2.1 Study Species

The red mangrove, *Rhizophora mangle* L. 1753 (Malpighiales, Rhizophoraceae), is an estuarine tree species present along the tropical and subtropical coasts of the Americas, eastern Africa, Bermuda, and a handful of outlying islands in the South Pacific (Tomlinson, 1986; Proffitt & Travis, 2014; DeYoe et al., 2020). *Rhizophora mangle* typically grows in the intertidal regions of sheltered coastlines, but can also be found in estuaries, tidal creeks, and occasionally along the edges of hypersaline salt pans (Duke, 2002; DeYoe et al., 2020). It is a dominant mangrove species across its range, including along peninsular Florida (DeYoe et al., 2020). Like other mangrove species, *R. mangle* functions as a foundation species by altering environmental conditions, providing nursery grounds for numerous fish species, and serving as a crucial primary producer within tropical and subtropical estuarine environments (Proffitt & Travis, 2005).

*Rhizophora mangle* is considered a self-compatible species (Nadia & Machado, 2014). Pollination in this species is mediated by both insects and wind (ambophilous pollen dispersal), which has been shown to effectively promote outcrossing and long-distance gene flow, but these outcrossing events are thought to be rare (Cerón-Souza et al., 2012). *Rhizophora mangle* produces viviparous propagules that mature for up to six months on maternal trees to lengths of 15-20cm (Goldberg & Heine, 2017; DeYoe et al., 2020). These propagules have considerable longevity at sea, surviving up to 3-4 months in the water column (Duke, 2002; Rabinowitz, 1978). However, propagules frequently recruit either directly underneath or nearby to maternal trees (Sengupta et al., 2005; Sousa et al., 2007; Goldberg & Heine, 2017) and maximum tidal action via king tides and major weather events is likely required to move propagules significant distances (Goldberg & Heine, 2017).

### 2.2 Field sampling

We sampled six populations of *R. mangle* between June 9 and June 26 of 2015, in the west coast of central Florida (USA) within the following county and state parks: Anclote Key Preserve State Park (AC), Fort De Soto Park (FD), Honeymoon Island State Park (HI), Upper Tampa Bay Conservation Park (UTB), Weedon Island Preserve (WI), and Werner-Boyce Salt Springs State Park (WB) (Figure 1). At each population, we collected leaf tissue and 20 propagules directly from each of 10 maternal trees separated by at least 10 m from each other to maximize the range of genetic variation sampled within each population (Albrecht et al., 2013). With this design, propagules from each maternal tree were at least half-siblings but they could be more closely related due to the high selfing rate of *R. mangle* in the study area (Proffitt & Travis 2005). We maintained leaf tissue of maternal trees on ice until transported to the Richards laboratory at the University of South Florida and then stored samples at −80 C (N=60). We refrigerated the propagules at 4° C for up to 14 days until we planted them in the greenhouse at the University of South Florida Botanical Gardens. In the greenhouse, propagules from four of the maternal trees at AC and nine of the maternal trees at FD failed to establish, so we returned to sample propagules and maternal tissue from 8 new maternal trees at FD on August 12 and 29, and from the same original maternal trees at AC on October 17.

**Figure 1:**
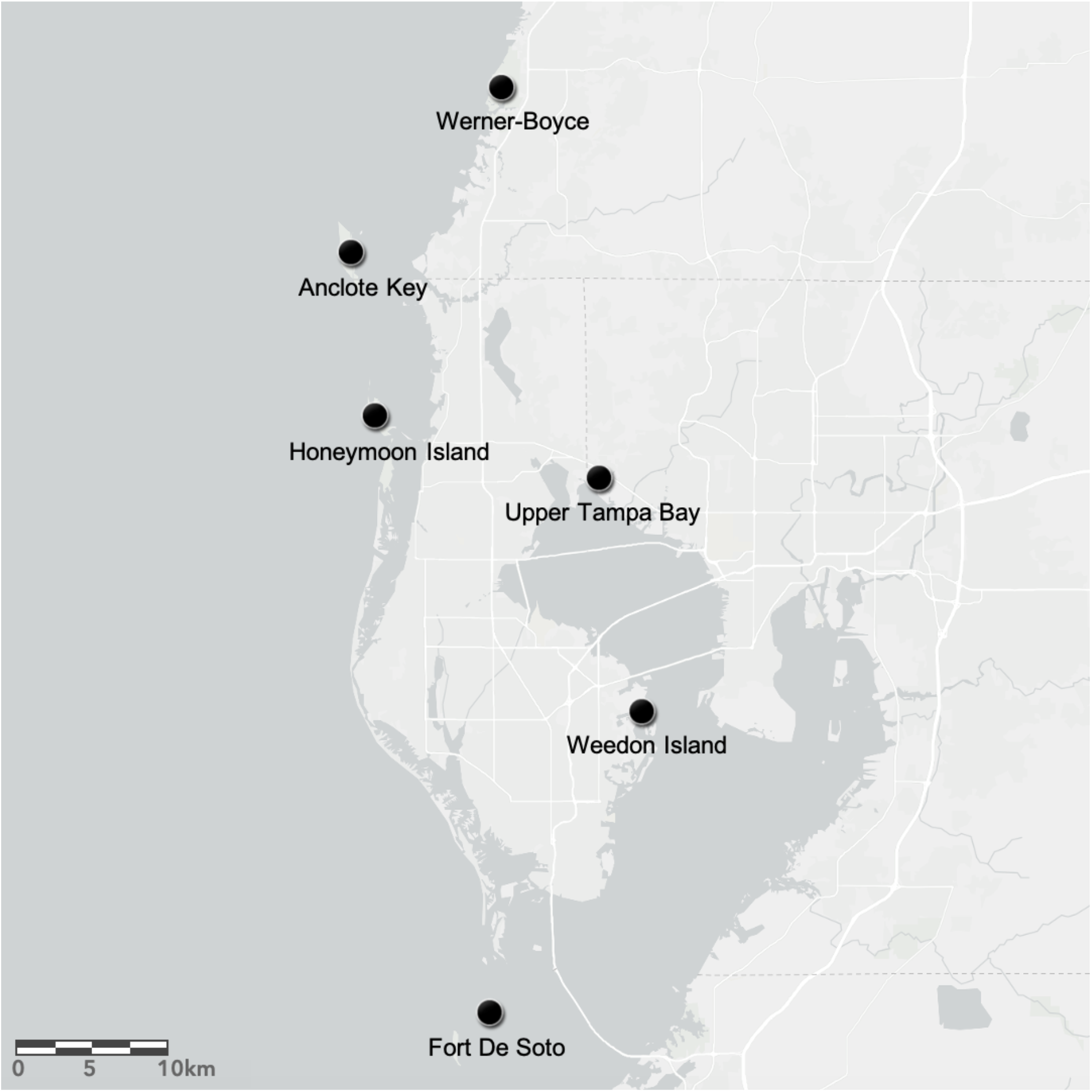
Map of six collection sites (aka populations) within the greater Tampa Bay region (FL, USA) generated in ArcGIS. We collected *Rhizophora mangle* leaves and propagules from ten maternal trees in Werner-Boyce Salt Springs State Park (WB), Anclote Key Preserve State Park (AC), Honeymoon Island State Park (HI), Upper Tampa Bay Conservation Park (UTB), Weedon Island Preserve (WI), and Fort De Soto Park (FD).

We planted propagules in 11.4 cm pots with a 50:50 sand and peat soil mixture and grew them for 9 months in the greenhouse at 18-29° C as part of a large common garden randomized block design experiment. We watered the plants daily with tap water until mid-October when we started applying salinity (15 ppt and 45 ppt reflecting the range of salinity measured in the field populations) and nutrient (no N amendment and high N, amended at approximately 3 mg N per pot each week, which is equivalent to a rate of 75 kg N per hectare per year) treatments twice per week in a full factorial randomized complete block design (N= 6 populations × 10 maternal families × 4 treatment combinations x 4-5 blocks x 1 replicate/block = 1150 plants, Langanke, 2017). Some families × treatment combinations were not represented in all five blocks due to limitations in the number of viable propagules. We harvested one block of plants per day between 2-7 May 2016, storing leaf tissue from each plant in paper envelopes, which we dried in a large glass container with silica gel (N=841 plants with leaves at the end of the experiment, ranging from 97-183 offspring per population). To assess genetic variation and structure, we chose 187 individuals representing 46 maternal families across the 6 populations (5-10 families per population). Since epigenetic variation can be induced by environmental variation, we selected plants from the low salinity, no nitrogen amendment treatment for the most part. We increased replication of some families for genetic (not epigenetic) diversity analyses with 29 plants that had received either high salt or high nitrogen treatments. By population, in the final group of samples that made it through the filtering process these 29 samples included AC (3 of 10 individuals), FD (5/47), HI (3/19), UTB (8/49), WB (4/24), WI (6/38).

### 2.3 Laboratory Methods

For genetic and epigenetic analyses, we isolated total genomic DNA from a total of 247 samples, including 60 maternal trees from the field and 187 offspring grown in the greenhouse. First, we disrupted approximately 80 mg of leaf tissue using stainless steel beads in a Qiagen TissueLyser II. Then, we extracted the DNA using the Qiagen DNeasy Plant Mini Kit following the manufacturer instructions with slight modifications that included an extended lysis step, a post-extraction clean-up with Buffer AW2, and elution in molecular grade water. The final concentration of DNA was quantified using the Qubit 3.0 Fluorometric dsDNA BR assay kit (Life Technologies).

We prepared libraries for epigenotyping-by-sequencing (epiGBS) following the methods outlined in van Gurp et al., (2016). In brief, we digested 400 ng of genomic DNA from each sample with the methylation-sensitive restriction enzyme PstI, and ligated methylated, non-phosphorylated barcoded adapters to the resulting fragments. We concentrated the libraries (NucleoSpin™ Gel and PCR Clean-up Kit), and size selected the fragments using 0.8x SPRI beads (Agencourt AMPure XP, Beckman coulter). We performed nick translation, bisulfite converted the fragments (EZ Lightning methylation kit, Zymo Research), and performed PCR amplification with the KAPA HIFI Uracil+ Hotstart Ready Mix (Roche). Finally, we quantified the libraries using the Qubit dsDNA assay kit, pooled them with equimolar concentrations (each sequenced library consisted of 96 multiplexed samples), and assessed their quality by analyzing 1 μl on a High Sensitivity DNA chip using an Agilent 2100 Bioanalyzer. We prepared libraries and sequenced paired-end reads of the 60 maternal plant samples and 36 randomly chosen offspring at the University of Florida Interdisciplinary Center for Biotechnology Research on one lane of the Illumina HiSeq 3000 (2 × 150bp) in February 2017. In August 2017, we prepared separate libraries for an additional 151 offspring and sequenced them at Novogene (HK) Company Limited in Hong Kong on two lanes of the Illumina HiSeq X-Ten System (2 × 150 bp): one lane contained 96 offspring samples, a second lane held 55 offspring samples along with 40 samples of another species prepared with the same protocol for another study (*Ceratodon purpureus*; Boquete, et al., unpublished).

### 2.4 Data processing

We processed the raw sequencing files using the pipeline provided by van Gurp et al., (2016) as in van Moorsel et al., (2019), available on https://github.com/thomasvangurp/epiGBS, with a bug-fix modification (https://github.com/MWSchmid/epiGBS_Nov_2017_fixed). Briefly, we demultiplexed, quality trimmed sequencing reads, and removed the barcode sequences, then used the processed reads for *de novo* reference construction. We mapped the reads to the *de novo* reference and called strand-specific single nucleotide polymorphisms (SNPs) and methylation polymorphisms (SMPs). *De novo* reference sequences were annotated with DIAMOND (protein coding genes; NCBI nonredundant proteins as reference; version 0.8.22; Buchfink, Xie, & Huson, 2015) and RepeatMasker (transposons and repeats; Embryophyta as reference species collection; version 4.0.6; Smit, Hubley, & Green, 2013– 2015). This annotation was used to classify the genetic variants (SNPs) and epigenetic variants (SMPs) into the different genomic features including genes, repeats, and transposons.

SNPs and SMPs were filtered to include only loci with a minimum coverage of 5 (i.e. 5 sequencing reads mapping to each locus) within each individual and across at least 5 individuals with a maximum coverage of 10,000 in at least five samples per population and type (maternal trees and offspring). Samples for which fewer than 60% of the SNP or SMP sites passed this filter were removed. This removed 59 samples from the original design, one maternal tree from FD, and 58 offspring spread across populations. The final design includes 59 maternal trees and 129 offspring (between 7 and 39 offspring per population from 3-10 maternal trees per population). Data were filtered again with the final design using the same criteria as before, resulting in 48,964 SMPs and 62,944 SNPs.

### 2.5 Data analysis

We separated each of the filtered SNP and SMP datasets into two distinct datasets comprising maternal trees and offspring respectively. Thus, all analyses were performed on the maternal trees and on the offspring datasets separately. We did not directly compare both datasets due to the fact that the resulting filtered data sets from the maternal trees and the offspring did not overlap for the most part, reflecting technical differences in sample storage between the maternal trees and the offspring (i.e. frozen vs. dry), and that their libraries were prepared and sequenced at different times. All the analyses were performed in R version 3.5.1 (R Core Team 2018).

#### 2.5.1 Genetic analyses

We calculated mean and standard deviation of observed gene diversity and observed heterozygosity per locus for each population in the maternal trees (N= 59 maternal trees from 6 populations x 9-10 maternal trees per population) and offspring datasets (N=129 offspring from 6 populations × 7 to 39 offspring per population) based on SNPs with no missing values (49,796 and 885 SNPs in maternal trees and offspring respectively) using the function basic.stats within the R package hierfstat (Goudet, 2005).

We tested for genetic differentiation within and among populations of *R. mangle* using several methods. With the maternal trees data, we tested for differentiation among populations with three different approaches. First, we used an analysis of molecular variance (AMOVA) within the function poppr.amova in poppr (Kamvar, Tabima, & Grünwald, 2014) and the model y ~ population. To test the significance of the model we ran a randomization test with 999 permutations on the output of the AMOVA using the function randtest from the ade4 package (Dray and Dufour, 2007). Second, we obtained overall F_st_ and pairwise F_st_ values using the functions wc and genet.dist respectively from the package hierfstat, and calculated the confidence intervals of the pairwise F_st_ values using the function boot.ppfst, from the same package, with 999 permutations to determine whether F_st_ values were significantly different from zero, i.e. to find evidence of significant population differentiation. Finally, we calculated the G-statistic using the function gstat.randtest with 999 simulations implemented in the package hierfstat. For this analysis, we subsampled 3,000 from 49,796 SNPs with no missing values for the maternal trees. Finally, to identify SNPs that could be under selection, we tested for outliers with bayescan (version 2.1, Fischer et al., 2011; Foll and Gaggiotti, 2008). SNPs were identified as significant if the FDR was below 0.05.

For the offspring data, we tested for differentiation among families (i.e. among maternal trees) within populations and among populations using only families with at least three members, and populations with more than one family (i.e. N= 90 offspring individuals across 24 families from 5 populations: 8 FD, 2 HI, 4 UTB, 3 WB, and 7 WI families). As with the maternal tree data set, we performed an analysis of molecular variance (AMOVA) with the model: y ~ population + family(population). We also completed overall and pairwise F_st_ as well as G-statistics analyses using all 3,786 SNPs with no missing values for the offspring dataset.

We quantified the relationship between genome-wide genetic variation and population of origin in the case of the maternal trees, and population and family in the case of the offspring (N=90), using redundancy analysis (RDA). RDA is an ordination technique that summarizes the main patterns of variation in the response matrix, i.e. the scaled allele frequency matrix created from the SNP data (obtained using the function scaleGen from adegenet with NA.method set to “mean”; Jombart, 2008), which can be explained by our explanatory variables, i.e. population (for the maternal trees) or population and family (for the offspring). We used the function rda implemented within the vegan package (Oksanen et al., 2017) to fit the following models:

1. maternal trees allele frequency matrix ~ population;
2. offspring allele frequency matrix ~ population + family.

We tested the significance of the variation explained by our explanatory variables using a Monte Carlo permutation test with 999 permutations and obtained adjusted R^2^ using the function RsquareAdj from the vegan package. We corrected p-values for multiple testing using the false discovery rate (“fdr”) method implemented with the p.adjust function in the base package of R.

#### 2.5.2 Epigenetic analyses

For both maternal trees (N=59) and offspring plants (N=90), we calculated the DNA methylation level at each SMP and individual sample as the number of reads mapping to one position showing evidence of methylation divided by the total number of reads mapping to that position.

We used a multivariate test for homogeneity of dispersions to estimate epigenetic diversity, i.e. variation in DNA methylation levels, for the maternal trees and offspring datasets following the approach of Anderson et al. (2006), which measures the average distance from each individual observation unit to their group centroid in a multivariate space using a dissimilarity measure. In line with this interpretation, we argue that the distance from each individual sample to its population centroid in a multivariate space generated using an epigenetic distance matrix provides an estimate of the extent of the variation in DNA methylation, i.e. epigenetic variation. Then, the average distance of each population can be compared to look for significant differences in the amount of epigenetic variation among populations. To do so, we generated pairwise epigenetic distance matrices for maternal trees and offspring by calculating the average difference in DNA methylation level across all cytosines between each pair of samples. Then, we used this matrix to calculate the distance between each individual sample and its population centroid using the function betadisper from the vegan package. We tested for differences in dispersion among populations using a permutation-based test of multivariate homogeneity of group dispersions on the output of betadisper with 9999 permutations. When this test was significant, we used the Tukey's Honest Significant Difference test to check which populations differed in their average distance to the centroid, i.e. in their levels of epigenetic variation. Finally, to compare genetic and epigenetic diversity levels, we used this approach to calculate the distance from each sample to its population centroid using genetic distance matrices. Genetic distances were calculated as the average distance of all per-SNP differences between two individuals. For each SNP, the distance was set to 0 if both alleles were identical, 1 if both alleles were different, and 0.5 if one allele was different.

We tested for differences in overall DNA methylation levels, i.e. the average percent DNA methylation per individual, and its standard deviation. We calculated average and standard deviation of percent DNA methylation for each separate sequence context (ie. CG, CHG and CHH) or across all sequence contexts, and then we used a general linear model (functions lm and anova) to test for significant differences among populations (maternal trees data) or among populations and families nested within populations (offspring data).

To assess the effect of population (for the maternal trees), and population and family (for the offspring) on genome-wide epigenetic variation, for each separate sequence context (i.e., CG, CHG and CHH) or across all sequence contexts, with and without accounting for their genetic structure, we used RDA and partial constrained RDA, respectively. Partial constrained RDA allows for “conditioning” the analysis of epigenetic variation with genetic data which we summarized with principal component analysis (PCA). The use of the “family” term in this analysis represents a composite of the maternal genetic and non-genetic contributions to the offspring epigenetic patterns since this term is not simply defined by maternal sequence patterns. Instead the “family” term is a categorical representation such that data for the propagules is explained by the association with the maternal tree more generally.

For the RDA, we used only SMPs with complete data, i.e. no missing values across samples: 41,164 (3,416 in CG, 10,432 in CHG, and 27,316 in CHH) for maternal trees and 9,038 SMPs (766 in CG, 2,549 in CHG, and 5,723 in CHH) for offspring. Similar to the genetic analyses, we only used families with at least three members, and populations with more than one family. First, we summarized the genetic data into principal components (PCs). We used the first 13 PCs for the maternal trees data which combined explained ~31% of the genetic variation in each of the three contexts. For the offspring, we used 12, 13 and 12 PCs for CG, CHG and CHH contexts respectively which explained 31, 30 and 31% of the variation respectively. Then, we ran the three following models to predict DNA methylation in the maternal trees:

1. maternal trees DNA methylation matrix ~ population;
2. maternal trees DNA methylation matrix ~ PCs from maternal trees genetic data;
3. maternal trees DNA methylation matrix ~ population + Condition(PCs from maternal trees genetic data).

We ran five similar models to predict DNA methylation in the offspring plants:

1. offspring DNA methylation matrix ~ population;
2. offspring DNA methylation matrix ~ family;
3. offspring DNA methylation matrix ~ PCs from offspring genetic data;
4. offspring DNA methylation matrix ~ population + Condition(PCs from offspring genetic data);
5. offspring DNA methylation matrix ~ family + Condition(PCs from offspring genetic data).

As for the genetic data, we tested the significance of the variation explained by our explanatory variables using a Monte Carlo permutation test and obtained adjusted R^2^, and adjusted p-values for multiple testing using the FDR method.

To test how much of the epigenetic (methylation) differentiation could be attributed to differences among populations, and how much of the epigenetic variation was associated with the populations after controlling for differences in sequence variation physically linked to the epigenetic variation, we modelled the average DNA methylation level of each 50-250 bp long fragment in response to the sequence context (CTXT), the population (POP) and its interaction with context (CTXT:POP), and the genotype of the fragment (GENO) and its interaction with context (CTXT:GENO) fitted in this order (percent methylation ~ CTXT + POP + CTXT:POP + GENO + CTXT:GENO). We then compared this result to an alternative model in which GENO and POP and their interactions with CTXT were switched (percent methylation ~ CTXT + GENO + CTXT:GENO + POP + CTXT:POP). We ran these models in R with the function anova() that uses type-I (i.e. sequential) tests.

Therefore, the first model tests for epigenetic differentiation between populations irrespective of the underlying sequence differences, and the second model tests whether there was epigenetic differentiation between populations that could not be explained by the underlying DNA sequence. For the offspring, we used similar models but further included the family term. We only used fragments which passed the coverage filters described above. Models were calculated with the functions lm and anova in R (version 3.6.1). Results from all reference sequences were collected and *P*-values for each term were adjusted for multiple testing by the FDR method (Benjamini & Hochberg, 1995). As noted previously (van Moorsel et al., 2019), this model is a good proxy for close-*cis* associations. However, given that it doesn't account for far-*cis* or *trans* associations, it tends to overestimate the proportion of epigenetic variation that is unlinked to genetic variation.

Finally, we identified differentially methylated cytosine positions (DMPs) between pairs of populations for the maternal trees and the offspring datasets using DSS (Feng, Conneely, & Wu, 2014) and adjusting for false discovery with FDR. This package models the DNA methylation level at each position within each group using a beta-binomial distribution with arcsine link function, and then performs Wald tests to detect differential methylation between groups at each position.

## 3. Results

### 3.1 Population genetics

We found overall low levels of genetic diversity among populations, with observed gene diversity values ranging between 0.009-0.012 and heterozygosity between 0.010-0.014 for the maternal trees, and 0.039-0.051 and 0.050-0.064 for the offspring (Table 1). We used three methods to examine genetic structure of the maternal trees, which all provided evidence of significant genetic differentiation among field populations of *R. mangle*. The randomization test performed on the output of the AMOVA was highly significant (Table 2), similar to the Monte Carlo permutation test carried out on the output of the RDA (Table 3), and the test on the significance of the G-statistic (G-stat = 39.9; p = 0.048). Yet, the amount of variation explained by the population of origin was rather low. According to the AMOVA, the bulk of the genetic variance is found within (99.4%) rather than among (0.63%) populations. Similarly, the RDA showed that population explains only 0.14% of the genetic variation. We found evidence for significant genetic differentiation between all population pairs except AC-HI and UTB-WI (Figure 2), the overall F_st_ was very low (0.003) and pairwise F_st_ values ranged between 0.0005 (UTB-WI) and 0.0081 (WB-WI; Figure 2).

**Table 1:** Mean and standard deviation (Std. Dev.) of observed gene diversity (Hs) and observed heterozygosity (Ho) per locus for each population in the maternal trees and offspring datasets calculated based on SNPs with no missing values. N: number of samples included in the analysis.

| Dataset | Population | N<br>(# families) | Observed gene diversity<br>(Hs) |  | Observed heterozygosity (Ho) |  |
| --- | --- | --- | --- | --- | --- | --- |
|  |  |  | Mean | Std. Dev. | Mean | Std. Dev. |
| Maternal trees | AC | 10 | 0.012 | 0.044 | 0.014 | 0.068 |
|  | FD | 9 | 0.011 | 0.045 | 0.013 | 0.067 |
|  | HI | 10 | 0.011 | 0.043 | 0.013 | 0.066 |
|  | UTB | 10 | 0.009 | 0.041 | 0.011 | 0.066 |
|  | WB | 10 | 0.010 | 0.041 | 0.011 | 0.065 |
|  | WI | 10 | 0.009 | 0.041 | 0.010 | 0.065 |
|  | Overall | 59 | 0.010 | 0.043 | 0.012 | 0.067 |
| Offspring | AC | 7 (3) | 0.048 | 0.111 | 0.056 | 0.155 |
|  | FD | 39 (10) | 0.050 | 0.098 | 0.064 | 0.169 |
|  | HI | 12 (6) | 0.047 | 0.103 | 0.055 | 0.148 |
|  | UTB | 25 (9) | 0.041 | 0.098 | 0.050 | 0.159 |
|  | WB | 16 (6) | 0.045 | 0.102 | 0.057 | 0.166 |
|  | WI | 30 (8) | 0.039 | 0.097 | 0.055 | 0.176 |
|  | Overall | 129 (42) | 0.045 | 0.102 | 0.056 | 0.162 |

**Table 2:** Analysis of molecular variance (AMOVA) carried out on the maternal trees and offspring datasets separately. Sigma: amount of genetic variance found among and within the predictor (population or family); Percent (%): percentage of genetic variance found among and within the predictor (population or family); Phi (ϕ): estimate of the extent of genetic differentiation among populations. ***: p<0.001; ns: not significant.

| | | Sigma | Percent (%) | Phi ($\phi$) |
| --- | --- | --- | --- | --- |
| Maternal trees | Among populations | 1.6789 | 0.629 | 0.0063*** |
|  | Within populations | 265.40 | 99.37 |  |
| Offspring | Among populations | 0.0684 | 0.016 | 0.0002<sup>ns</sup> |
|  | Among families | 4.4994 | 1.026 | 0.0103*** |
|  | Within families | 433.95 | 98.96 | 0.0104*** |

**Table 3:**
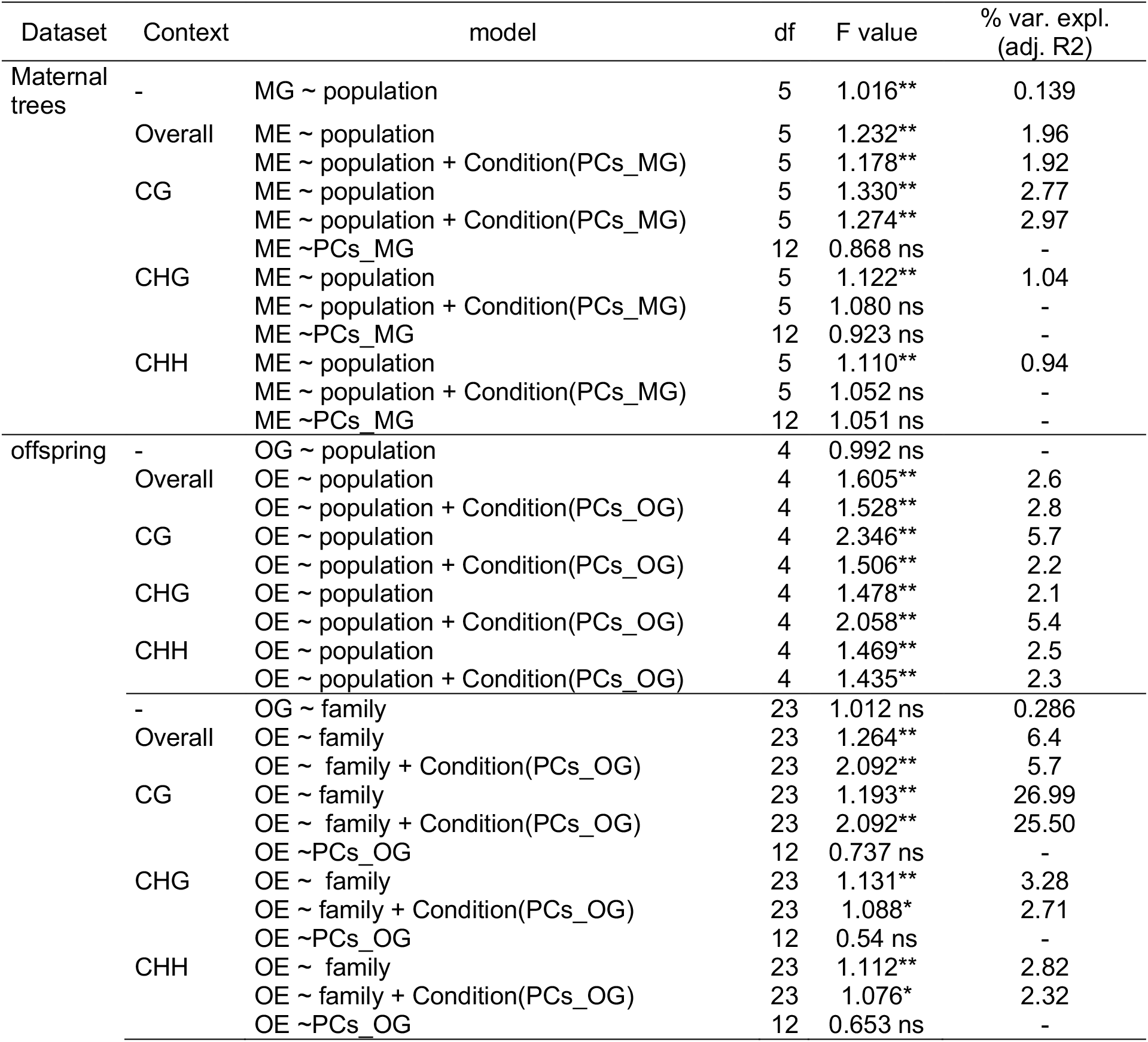
Results of the redundancy analysis (RDA) showing the percentage of genetic and epigenetic variance explained by population (in maternal trees data set), and population and family (in offspring data set) with and without adjusting for the variance explained by the genetic component. The output of the Monte Carlo permutation test (F value and significance) is also shown. Context: sequence context for DNA methylation; df: degrees of freedom. % var. expl. (adj. R^2^): percent of variance explained as the R^2^ adjusted for multiple comparisons; MG: maternal trees genetic matrix; ME: maternal trees epigenetic matrix; PCs_MG: matrix of principal components summarizing the maternal trees genetics; PCs_OG: matrix of principal components summarizing the offspring’s genetics; **: p<0.01, *: p<0.05, ns: not significant.

**Figure 2:**
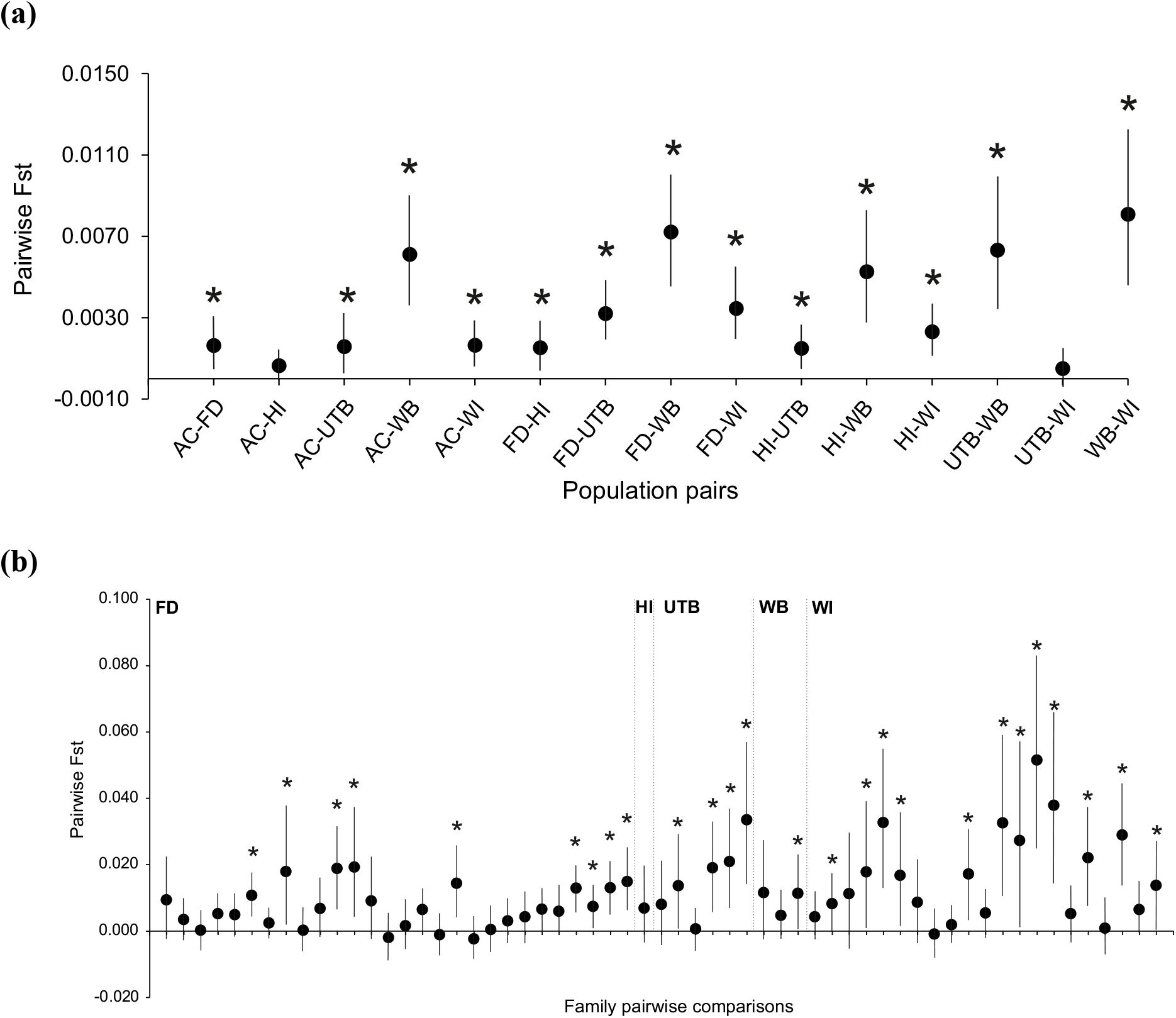
Pairwise F_st_ values between field populations of *Rhizophora mangle* in the maternal tree data (a), and between families within populations in the offspring data (b). Bars correspond to the 2.75 and 97.5% confidence intervals obtained using bootstrapping. Stars at the top of the graph highlight significantly genetically differentiated population pairs, i.e. F_st_ values different from 0.

We visualized the genetic data by means of PCA using the complete SNP dataset as well as the 5% most differentiated SNPs, finding that a clear separation among populations was only possible when using the 5% most differentiated loci (Figure 3). We found that the significant genetic differentiation among maternal trees of *R. mangle* yielded by our statistical analyses was principally due to the distinctness of WB, and possibly of HI, from the rest of the populations. The separation of WB from all other populations was also reflected in the higher pairwise F_st_ values between WB and the others (Figure 2). Finally, our analysis yielded 277 SNPs showing significant signs of differences among maternal trees of *R. mangle* collected in the field. The 277 SNPs were located in 111 different sequence fragments, out of which 26 had a high sequence similarity to known genes (descriptions in Table S2).

**Figure 3:**
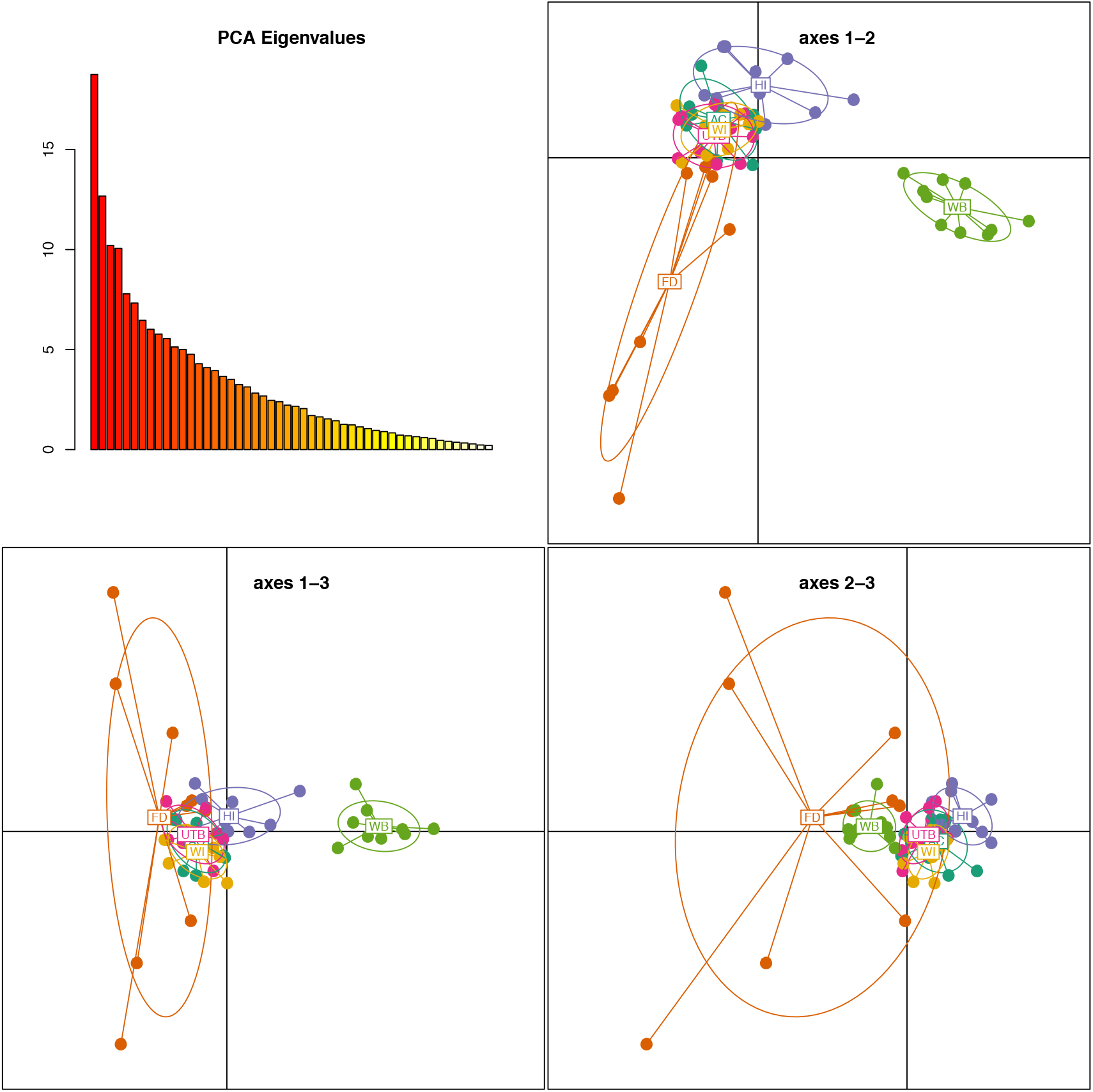
Visualization of the genetic structure of the maternal trees of *Rhizophora mangle* using only the 5% of the most differentiated SNPs.

Results of the genetic analyses on the offspring are similar to that found for the maternal trees; the AMOVA showed significant genetic differentiation among families but this predictor explained only 1% of the genetic variance. The majority of the variance was found within families (99%) and population did not significantly explain any proportion of the genetic variation of the offspring (Table 2). On the other hand, the RDA model with population and family did not explain any of the variation of the offspring genetics (Table 3). The G-tests for differentiation between families within populations were significant in two out of five tested populations (WI: G-stat = 171.6, p = 0.001, UTB: G-stat = 131.5, p = 0.005). Again, the overall F_st_ value was very low (0.022) and pairwise F_st_ ranged between −0.0023 and 0.0515 (Figure 2).

### 3.2 Population epigenetics

DNA methylation across all contexts was around 9% for all populations in the maternal trees dataset whereas for the offspring this value ranged between 11-17% (Table 4). Similarly, DNA methylation levels in CG, CHG and CHH contexts were close to 28, 23, and 1% respectively for all populations in the maternal trees dataset and slightly higher in the offspring (29-33%, 25-30%, and 4-9% for CG, CHG and CHH contexts respectively). The average distances from each sample to its population centroid estimated as a proxy of the amount of epigenetic variation range between 0.02 and 0.03 in the maternal trees and between 0.05 and 0.08 in the offspring (Figure 4). The tests for homogeneity of multivariate dispersions were significant for both the maternal trees and the offspring datasets (F = 23.5, p < 0.001; F = 16.0, p < 0.001 respectively) revealing significant differences in the levels of epigenetic diversity among populations in both datasets. The multiple pairwise comparisons within each dataset showed that these differences were due to the greater epigenetic diversity found in WB in the maternal trees. In the offspring, FD, HI and UTB showed higher levels of epigenetic diversity than UTB and WI (Figure 4). The average distances to centroid estimated with the genetic data were an order of magnitude lower for the mothers (ranging between 0.007 and 0.01), and between 2.5x and 4x times lower for the offspring (data not shown).

**Table 4:** Average DNA methylation levels for each context and across all contexts for each population for maternal tree and offspring separately.

| Dataset | Population | CG | CHG | CHH | ALL |
| --- | --- | --- | --- | --- | --- |
| Maternal trees | AC | 27.6 | 23.2 | 0.7 | 8.7 |
|  | FD | 27.9 | 23.4 | 0.9 | 8.9 |
|  | HI | 27.7 | 23.3 | 0.8 | 8.7 |
|  | UTB | 27.6 | 23.3 | 0.9 | 8.8 |
|  | WB | 28.2 | 23.5 | 1.1 | 9.1 |
|  | WI | 27.7 | 23.2 | 0.7 | 8.7 |
| Offspring | FD | 33.2 | 29.1 | 8.8 | 16.0 |
|  | HI | 33.5 | 29.5 | 9.5 | 16.6 |
|  | UTB | 30.1 | 25.9 | 4.5 | 12.1 |
|  | WB | 32.0 | 27.9 | 7.3 | 14.6 |
|  | WI | 29.4 | 25.4 | 3.8 | 11.4 |

**Figure 4:**
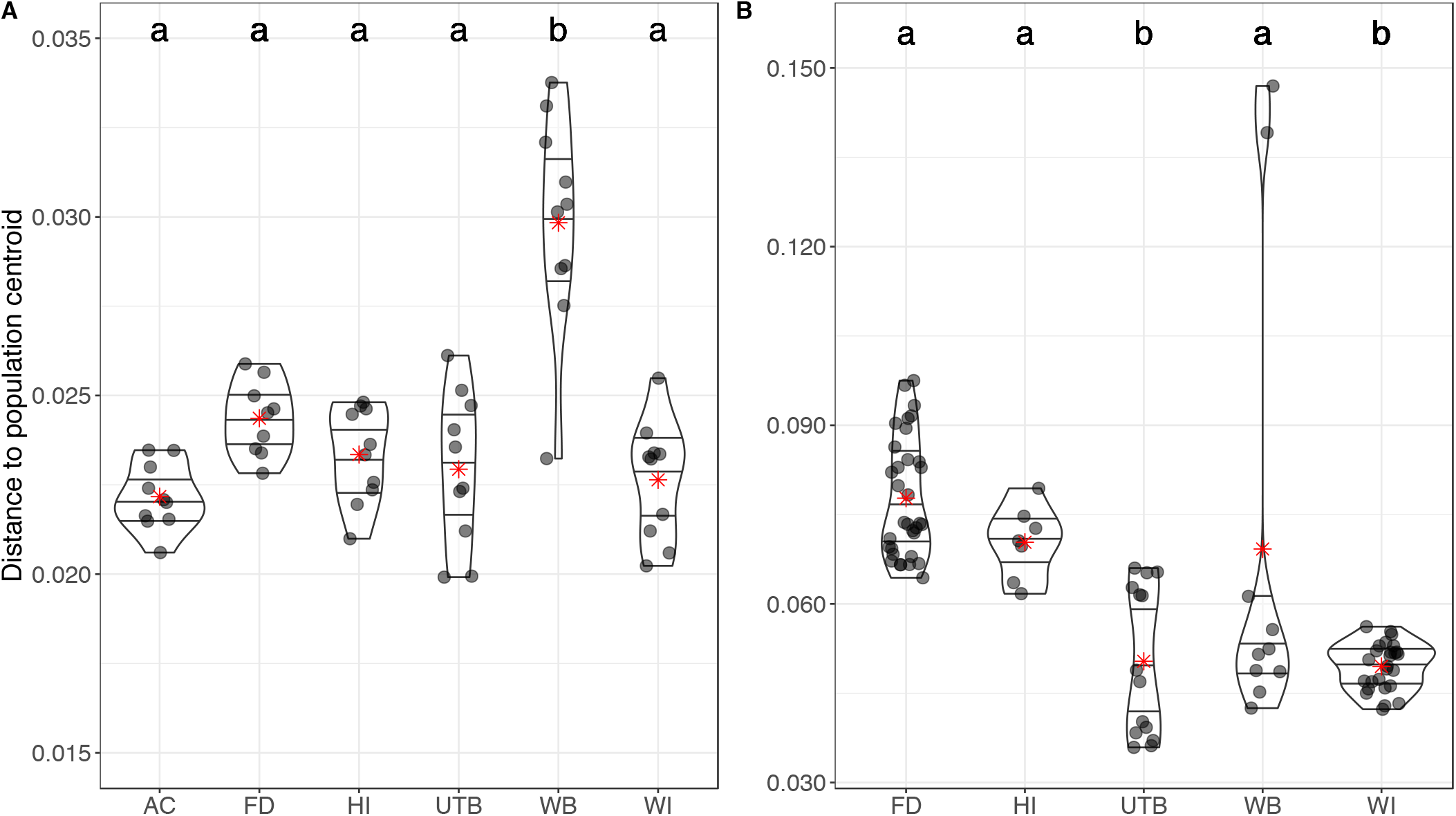
Distance from each individual sample to its corresponding population centroid calculated using epigenetic distance matrices for the maternal trees (A) and offspring datasets (B). Lines within the violin plots mark the 25, 50, and 75% quartiles of the distribution; letters inside the graphs summarize the results of the multiple pairwise comparisons where populations sharing letters do not differ significantly in epigenetic diversity; red stars: average distance to centroid for each population.

The linear models that test for differences in average DNA methylation levels and standard deviation in DNA methylation showed that population of origin significantly explains 75% of the variation in average and 52% of the variation in standard deviation for the maternal trees if data from all sequence contexts were used. Within individual contexts, the numbers were similar but there was no significant association between population of origin and average DNA methylation in the CG context (Table S1). Among the offspring, family alone significantly explained 79% of the variation in average and 74% of the variation in standard deviation if data from all sequence contexts were used. At least half could be attributed to differences between population (47% of the variation in average and 37% of the variation in standard deviation). Within individual sequence contexts, the results were similar for the average DNA methylation levels. However, the association between population of origin and variation in standard deviation in CG and CHG context were not significant (Table S1).

The RDA analysis on the effect of population (for the maternal trees), and population and family (for the offspring) on genome-wide epigenetic variation showed that epigenetic variation is significantly structured in both datasets. Population significantly explained 1.96 of the total epigenetic variation across all sequence contexts in the maternal trees and 2.6% of the epigenetic variation in offspring; family explained 6.4% of the total epigenetic variation across all sequence contexts in the offspring. Additionally, population explained 2.77, 1.04, and 0.94% of the epigenetic variation in CG, CHG and CHH respectively for the maternal trees (Table 3). For the offspring, family explained again a greater percent of epigenetic variation with 26.99, 3.28, and 2.82% in CG, CHG and CHH respectively against the 5.7, 2.1, and 2.5% in CG, CHG and CHH respectively explained by population (Table 3).

The partial RDA in which the same models were conditioned on the maternal and offspring genetic data (based on PCs) showed similar results for the effect of population in maternal trees (explaining 1.92% of the variation) and offspring (2.8%), and of the family term in the offspring (5.7%; Table 3) across all sequence contexts. However, when examining each context separately, the effect of population only remains significant in the CG context for the maternal trees and explains 2.97% of the variation (Table 3). For the offspring, both population and family remain significant in all contexts after accounting for the offspring’s genetic component; population explains similar levels of the variation in CG, CHG and CHH contexts respectively, while family explains 25.5, 2.7, and 2.3% of the variation in CG, CHG and CHH contexts respectively. The genetic component does not significantly explain any of the epigenetic variation in either the maternal or the offspring data or across all contexts and each sequence context separately (Table 3).

The differential methylation analysis with DSS comparing DNA methylation levels at individual cytosines between pairs of populations yielded between 0.02 and 1.1% significant cytosines in the maternal trees and between 0.1 and 4.5% significant cytosines in the offspring. Gene annotations of these DMPs are shown in Tables S3 and S4. For maternal trees, the most pronounced differences were found between WB vs. WI (1.13% significant Cs) and between WB vs. UTB (1.08% significant Cs). Almost no differences were found between AC vs. FD, HI, UTB, and between UTB vs. FD, HI, WI (≤0.05% significant Cs). In the offspring the higher number of significant Cs were found between WI vs. FD (4.5%), WI vs. HI (2.7%), and UTB vs. HI (1.8%). The smallest differences in the offspring were found between UTB vs. FD, WB, WI and between FD vs. WB (between 0.1% and 0.8% significant Cs). Comparing family pairs in the offspring resulted in between 0.4 and 14.7% significant Cs. On average, family pairs differed significantly in 3.3% of all cytosines. The greatest differences were found between family WB6 and all other 23 families (between 10.2 and 14.7% significant Cs; Table S5).

We detected individual fragments in which the epigenetic variation was unlinked to sequence variation on the same reference fragment (*i.e.*, in close-*cis*). In the maternal trees, we found that population and its interaction with the sequence context (POP & CTXT:POP) could significantly explain differences in DNA methylation in 19.3% of all fragments (FDR < 0.05). However, if the terms testing for population were fitted after the factor accounting for the sequence of the fragments (GENO & CTXT:GENO), only 5.9% of all fragments were still significant for POP & CTXT:POP indicating that differences in these fragments could not be explained by the underlying sequence differences in close-*cis*. In the offspring, POP & CTXT:POP was significant for 82.7% of all fragments if fitted first. In addition, terms testing for differences between families (MOTHER & CTXT:MOTHER) were also significant for 68.2% of all fragments. Notably, even if GENO & CTXT:GENO was fitted first, POP & CTXT:POP and MOTHER & CTXT:MOTHER were significant in 60.5 and 45.8% of all fragments, respectively.

## 4. Discussion

Conservation biologists strive to preserve biodiversity and face the enduring challenge of doing so in the context of changing environmental conditions. While the capacity to respond to environmental challenges ultimately relies on phenotypic variation, deciphering the mechanisms that contribute to phenotypic variation is a challenging task that requires a better understanding of the complex interactions of genetic and non-genetic mechanisms. DNA methylation has been associated with regulation of gene expression (and therefore phenotype) in some contexts, and has been proposed to contribute to phenotypic variation, particularly in populations with low genetic diversity (Verhoeven & Preite 2014; Douhovnikoff & Dodd 2015; Richards et al., 2017; Xie et al., 2019; Mounger et al., 2020). Investigating biodiversity at these different molecular levels can contribute to our understanding of response in foundation species like mangroves, which inhabit dynamic coastal landscapes and are constantly under threat from various anthropogenic challenges.

Populations of *R. mangle* around Tampa Bay are near the species northern limit, dictated largely by periodic freezing events (Kennedy et al., 2016). In addition, they could be more vulnerable to changing conditions due to increased isolation and reduced genetic diversity (Polidoro et al., 2010; Sandoval-Castro et al., 2012; Kennedy et al., 2016), resulting from inbreeding, limitations in dispersal ability, and increased environmental pressures (Sandoval-Castro et al., 2012). Our study confirmed that these populations had low genetic diversity, but we also found that differences among populations explained very little of the variation. On the other hand, there was considerable epigenetic variation and more of the epigenetic variation was explained by differences among populations for both the maternal trees and offspring, while maternal family explained the largest percentage of the variation in epigenetic variation in the offspring plants. This pattern of DNA methylation in the offspring plants suggests that propagules maintain some level of epigenetic variation inherited from the maternal plant or maternal environment even when they are grown under common garden conditions, which could have important implications for how these propagules can respond to environmental challenges.

### 4.1 Red mangrove population genetics

While high levels of diversity in both heterozygosity and allelic number have been reported from populations of *R. mangle* along the Pacific coast of Nicaragua (Bruschi et al., 2014), and Colombia (Arbelaez-Cortes et al., 2007), Pil et al., (2011) compared these findings to populations of *R. mangle* along the Brazilian coast and determined that genetic diversity was lower in Brazil. They also found considerable genetic structuring between the northern and southern Brazilian populations, possibly resulting from the last glacial period (Pil et al., 2011). Studies at the current range edge have also reported much lower levels of diversity (Polidoro et al., 2010; Sandoval-Castro et al., 2012; Kennedy et al., 2016). In our study, we found overall low levels of diversity, and that most of the genetic variation was found within populations and even more so within families. This type of genetic structure follows from the known levels of inbreeding of the species followed by mixing of the populations through the dispersal of propagules (Pil et al., 2011; Francisco et al., 2018). Although differences among populations explained very little of the genetic variation, almost all pairwise comparisons showed significant fine scale genetic differentiation except for the Anclote Key (AC) to Honeymoon Island (HI) and Upper Tampa Bay (UTB) to Weedon Island (WI) comparisons. The lack of differentiation specifically between these pairs of populations might be explained by spatial proximity and propagule dispersal. UTB and WI are the only two populations sampled that are within the mouth of the bay. AC and HI are both barrier islands that are geographically close to one another and therefore have a conceivably greater chance for dispersal between these two islands than between other populations (Figure 1).

A study by Albrecht et al., (2013) provides insight for interpreting our findings in the context of the larger range of the species, since they compared genetic diversity among Florida and Caribbean populations. They found high genetic structuring among the populations, and that populations from the Gulf Coast of Florida had much higher structuring compared to those along the Atlantic Coast suggesting that there is limited gene flow along the Gulf Coast and across to other parts of the species range, including the Caribbean islands and throughout Florida (Albrecht et al., 2013). They suggest that genetic structuring and loss of genetic diversity in some populations are related to habitat loss via human development (e.g. the Atlantic Coast of Florida has experienced more extensive habitat loss than the Gulf Coast). While our findings of minimal genetic diversity among populations run contrary to those in Albrect et al. (2013), this could in part be explained by significant urbanization and resultant habitat loss in the Tampa Bay region.

Our findings of limited genetic variation in *R. mangle* are similar to other studies in this part of the species range, but contrast with several other foundation coastal species of the southeastern U.S.A., which are outcrossing grasses or rushes that exhibit much higher levels of genetic diversity. Studies on native southeastern U.S. *Spartina alterniflora* populations have reported diversity levels that are comparable to other outcrossing grasses, despite the fact that this species also spreads prolifically by clonal reproduction (Richards et al., 2004; Foust et al., 2016, Robertson & Richards, 2017). Tumas et al., (2019) found greater genetic diversity in Gulf of Mexico than Atlantic coast populations of the salt marsh foundation plant *Juncus roemerianus*, but like in *R. mangle*, measures of genetic diversity varied dramatically across the range. The authors suggest this could be the result of differences in plant community and disturbance regimes or reflect a relationship with population size.

### 4.2 Population epigenetics

The limited genetic diversity in these populations of *R. mangle* might be cause for concern considering the important ecosystem functions provided by this foundation species, but what really matters is how the species can maintain phenotypic response to challenging environments. Like in several other studies of coastal foundation species, we found epigenetic variation was high in *R. mangle* (based on test for dispersion; see also Lira-Medeiros et al., 2010; Foust et al., 2016; Robertson et al., 2017; Alvarez et al., 2020). Further, this variation was significantly associated with population for both maternal trees and offspring plants, and even more significantly associated with family for the offspring plants. Although, using a categorical family term in the analysis does not allow for prescribing effects specifically to the mother’s genetic, epigenetic, or other non-genetic contributions to the offspring epigenetic matrix, the family term does represent a holistic contribution from the maternal tree to offspring and in our study explains the largest portion of the variation (approximately 6% overall and 25% of the variation in the CG context). This provides some of the first evidence for epigenetic inheritance in a coastal foundation species.

While it has been established that genetic variation can have considerable effects on epigenetic variation (Becker et al., 2011; Dubin et al., 2015; Sasaki et al., 2019), we found significant epigenetic structure in both maternal trees and offspring that could not be explained by the genetic sequence (i.e. genetic variation in close *cis*) of the fragments. Instead, population of origin explained more of the variation in DNA methylation than for sequence variation. This finding was true not only in the field collected plants, but also in the propagules grown in a common garden. Several other studies have found that epigenetic patterns that are associated with habitat can persist in common gardens, suggesting that environmentally induced epigenetic differences can be inherited, and contribute to diversity (Richards et al., 2012; Xie et al., 2015; Robertson et al., 2020). However, in our study, we collected the propagules from the field and they had already matured on the maternal plants. Therefore, some important early developmental responses would reflect the maternal environment. Further study is required to truly control for environmental, maternal, and genetic effects which may not be possible in such a long-lived tree species.

These findings suggest that epigenetic variation could contribute to heritable differences in *R. mangle*, but this would depend also on which propagules survive the various stages of selection before establishment in the field. A recent study of propagule recruitment in *R. mangle* at its range edge near Jacksonville, Florida found that just two maternal trees contributed 79% of propagules that reached branching stage. Propagule survival was higher in populations within the range core compared to the range edge, even though there was a longer propagule development period and greater reproductive output among trees at the range edge (Goldberg & Heine, 2017). So far, very little is known about how this or any coastal foundation species survives the different selection pressures across the various stages of establishment and spread. Variation in these selection pressures will be amplified by the pressures attendant to anthropogenic climate change.

## 5. Conclusions

The field of conservation biology relies on identifying the capacity of organisms to respond to environmental challenges which ultimately relies on the manifestation of phenotypic variation through complex interactions of genetic and non-genetic mechanisms. We know that documenting the levels and structure of genetic variation is one piece of information that is important for conservation, but how that information is translated into function largely remains an enigma. We have provided another piece of the puzzle for the coastal foundation plant *Rhizophora mangle* that epigenetic variation (namely DNA methylation) is inherited and could be an important component of diversity for this species. However, our interpretation of how this variation might be involved is limited due to the small portion of the genome sample with our RRBS approach and the limited genomic resources (see also van Moorsel et al., 2019; Alvarez et al., 2020; Robertson et al., 2020). We look forward to the future of integrating novel molecular tools that can probe more deeply into the molecular underpinnings of response, as they will help shed light on the processes of development in the context of climate change.

## Supporting information

RM_epiGBS_SI

## Acknowledgments

We thank Samantha Blonder, Jordan Dollbaum, Maria Nikolopoulos, Shane Palmer, Bradley Biega, Bryan Lotici, Harper Cassidy, Jelena Dosen, Nancy Sheridan, and Dawei Tang for help with field collections, and with the experimental offspring plants in the greenhouse. We thank Stephen Savage and Priscila Magalhães Galdino for help with the epiGBS laboratory work. We thank the Coordenação de Aperfeiçoamento de Pessoal de Nível Superior (CAPES, 072/2014) – National Science Foundations (NSF, I-REU) bilateral grant for supporting the international research exchange between USF and the Jardim Botânico do Rio de Janeiro in Brazil for JM, SV, MA and RG. This work was supported by funding from the National Science Foundation (U.S.A.) DEB-1419960 (to CLR), Deutscher Akademische Austauschdienst (DAAD; MOPGA Project ID 306055 to CLR) and the European Union’s Horizon 2020 research and innovation programme under the Marie Skłodowska-Curie grant agreement No 704141-BryOmics (to MTB).

## Author contributions

CLR, CFL, JM, SV, and MHR conceived the study. CLR, JM, MTB, MWS, GAF, and DBL designed the experiments and analyses. JM, KL, SV, MA, and MHR collected plants and maintained experiments. JM, MHR, MTB, RG, and CAMW did the epiGBS laboratory work. MTB, MWS, and AWS analyzed the epiGBS data. CLR, JM, MTB, and MWS wrote the first draft of the manuscript. All co-authors provided input and revisions to the manuscript.

## Data accessibility

The pipeline scripts we used for this study are available at: https://github.com/thomasvangurp/epiGBS, with a bug-fix modification (https://github.com/MWSchmid/epiGBS_Nov_2017_fixed). The epiGBS sample design file, barcode file, pipeline output files (methylation.bed, consensus cluster, and vcf file), and annotation files are available on dryad. Raw sequence data files for epiGBS (Illumina reads after demultiplexing) will be submitted to NCBI. Links to this data will be made available in the “readme mangroves 2020” file on Dryad: https://doi.org/10.5061/dryad.3bk3j9khf

## Supporting Information

Supplemental tables are submitted as tables S1-S4. Table S5 is too large to submit on the manuscript submission site and is available here: https://www.icloud.com/iclouddrive/06gMNwnNaIBsEQ-MpT-_h873g#S5%5FDSSwithGeneAnnotation.offspringFams

## Supplementary Tables (provided on-line at the ManscriptOne for review)

Table S1: Analysis of variance of average DNA methylation and standard deviation of DNA methylation for each separate sequence context (ie. CG, CHG and CHH) or across all sequence contexts. For the offspring data, “Population” was tested against “Mother” (nested design or mixed model with Mother as random term).

Table S2: Genes mapped by fragments with SNPs showing signs of selection (sequence and description retrieved from the NCBI non-redundant protein database). From 111 fragments, 26 matched to genes. Note that one fragment may map to multiple genes.

Table S3: Differential cytosine methylation between populations using the mother data set. The first three columns fragment number (“chr”), the position within the fragment (“pos”), and the sequence context (“context”). Columns with the pattern FDR_<X>_vs_<Y> contain false discovery rates of a test comparing population X with population Y. Average DNA methylation levels for each population are given in the columns “AC”, “FD”, “HI”, “UTB”, “WB”, and “WI”. The remaining columns contain the annotation of the fragment, for example whether it matches to a gene and if yes, the gene name ID and description are provided.

Table S4: As S3 but using the offspring data set.

Table S5: As S4 but comparing the families instead of the populations (available here: https://www.icloud.com/iclouddrive/06gMNwnNaIBsEQ-MpT-_h873g#S5%5FDSSwithGeneAnnotation.offspringFams)

